# Mycol: A user-friendly app for automating analysis of microscopy images

**DOI:** 10.64898/2026.06.02.729113

**Authors:** Samuel Alan Bradley, Henry Webel, Giovanni Schiesaro, Marcel Skumantz, Olga Novillo-Sanjuan, Alexandra Panagou, Rafael Lucena-Marín, Emil D. Jensen, Antonio Di Pietro, Carlos G. Acevedo-Rocha

## Abstract

Microscopy image analysis is central to modern biology, yet many available platforms remain inaccessible to non-specialist users because they require advanced technical expertise, code-based workflows, extensive setup, or paid access. This creates a barrier for researchers who need reliable and fast image quantification but lack dedicated computational support. Here, we introduce Mycol, an open-source, machine-learning-assisted image analysis platform designed to be accessible and run on standard laptops with minimal setup. Mycol supports end-to-end workflows in which users annotate microscopy images, perform human-in-the-loop fine-tuning of machine learning models for automated segmentation and classification, deploy machine learning models, quality control predictions and quantitatively compare morphological and class frequency descriptors through a single intuitive interface. By combining machine-learning analysis with efficient quality control by humans, Mycol makes rapid and high-quality image quantification available to biologists without requiring specialist training. We demonstrate the utility of Mycol in diverse workflows using two economically important organisms, the crop pathogen *(Fusarium oxysporum)* and the blue mussel *(Mytilus edulis)*. Through Mycol, curated training sets were generated and high quality segmentation and classification models were obtained in each case. Deploying these models through Mycol decreased the time requirements and increased traceability of established cell counting workflows and facilitated a quantitative comparison of morphological parameters that reveals new patterns in early *M. edulis* larval development.

## Introduction

Converting microscopy images into quantitative datasets is a common task in biological research that, without computational assistance, is repetitive, time consuming and error-prone. Several specialist image-analysis platforms exist and have been recently reviewed (Liu et al. 2024), including ImageJ, Cellpose, CellProfiler, and Napari. ImageJ is a long-standing open-source platform for microscopy image analysis. Cellpose is a deep learning–based, general-purpose segmentation tool for 2D and 3D microscopy images, widely used for training custom models for segmenting cells and nuclei across diverse imaging modalities (Pachitariu and Stringer 2022; Stringer and Pachitariu 2025). CellProfiler is an open-source, modular platform widely used for high-throughput cell image analysis, supporting segmentation, object classification, and morphological feature extraction (Carpenter et al. 2006; Stirling et al. 2021). Napari is an interactive Python-based image viewer commonly used as a foundation for segmentation workflows and plugins, including Cellpose integration via cellpose-napari (Chiu et al. 2022).

While these platforms are powerful, their breadth of functionality often requires additional plugins and familiarity with complex interfaces, creating a barrier to entry for non-specialists. The Cellpose graphical user interface (GUI) offers a simpler interface but it is limited to segmentation and does not support classification or morphological analysis. As a result, many users whose primary focus is not image analysis face unnecessary complexity for relatively routine tasks. There is therefore a clear need for an integrated, streamlined solution that allows users to quickly train and apply machine learning models, perform quality control, and export concise, interpretable reports.

To address this gap, we developed Mycol: a streamlined, intuitive, and installation-friendly application for training Cellpose segmentation (Pachitariu and Stringer 2022) and DenseNet121 classification models (Huang et al. 2017), quality-controlling automated analyses, and comparing morphological descriptors of identified cells (**Fig. 1a**). Mycol is designed to lower the technical barrier to applying machine learning (ML) models in cell image analysis, enabling biologists to combine manual annotation with automated segmentation and classification within a single intuitive interface. We demonstrate, using two case studies in different species (*Fusarium oxysporum* and *Mytilus edulis*) and different technical set ups, that Mycol can improve the time efficiency and reporting clarity of existing protocols as well as facilitate new biological insights. Mycol is available for local installation (https://github.com/biosustain/mycol) to provide a low-barrier entry point to ML-assisted microscopy analysis for non-expert users with small-scale or occasional imaging needs.

**Figure 1.**
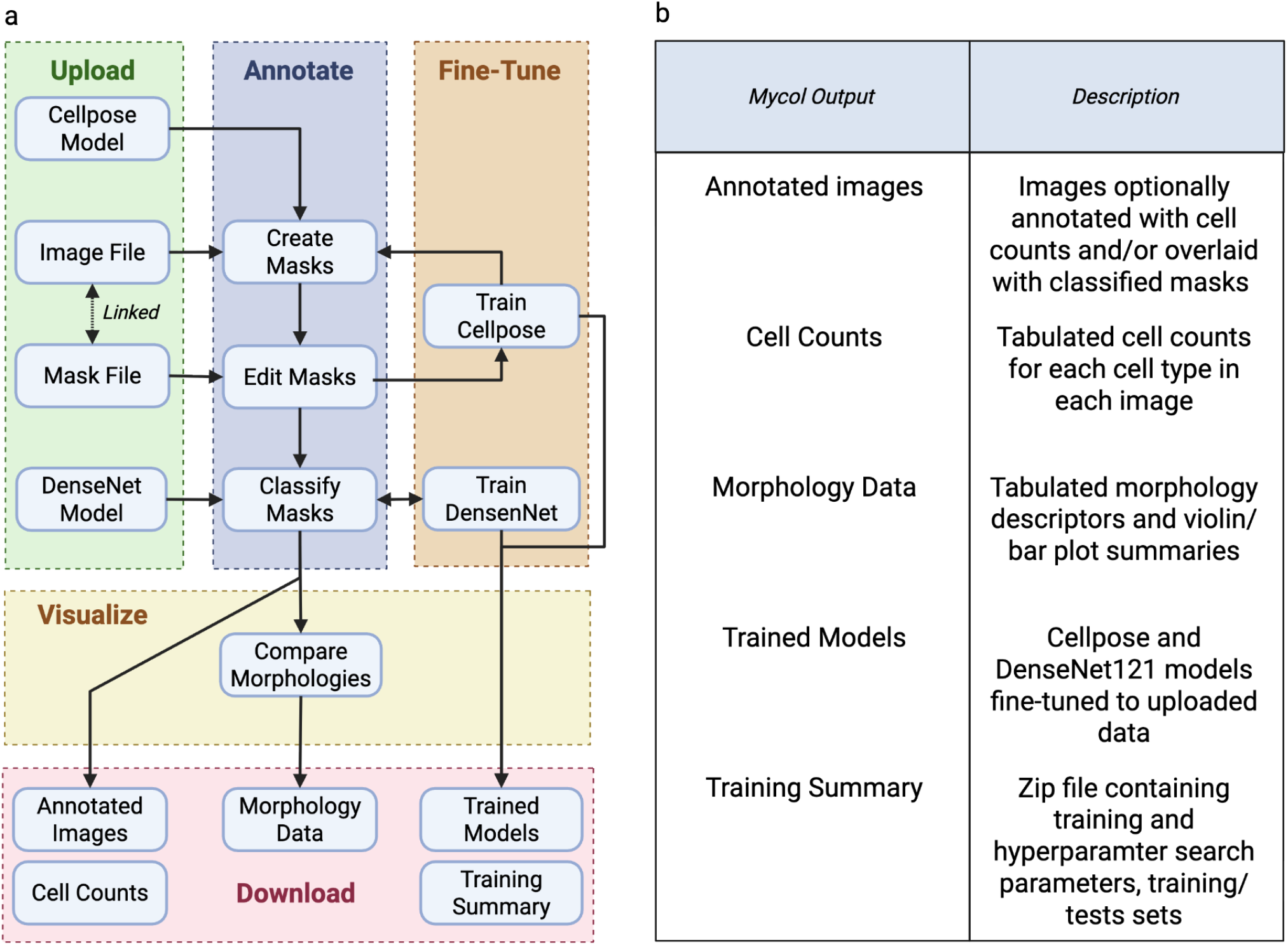
Mycol workflow. **a**) Schematic demonstrating the five modules in the Mycol application and how they fit together into a workflow. **b**) Brief description of Mycol outputs.

### Mycol Workflow: Upload Data and Models

All workflows begin with the *Upload Data and Models* page (**Supp. Fig. 1**), where users can import images (which can be from a microscope or even a smartphone), optional mask files associated with an uploaded image and previously trained Cellpose (segmentation) and DenseNet (classification) models (Huang et al. 2017; Pachitariu and Stringer 2022; Stringer and Pachitariu 2025; Mansourvar et al. 2024).

### Mycol Workflow: Creating a dataset of annotated images

The *Annotate Images* page integrates manual and ML-assisted tools for generating high-quality annotations (**Supp. Fig. 2**). Mycol provides an interactive image and two simple control panels that allow the user to easily add segmentation and classification annotations to their image dataset.

#### Segmentation Panel

users can draw or delete masks manually on an interactive image, guide segmentation using SAM2-based bounding boxes (Ravi et al. 2024), or apply fine-tuned Cellpose models in single-image or batch mode (Pachitariu and Stringer 2022; Stringer and Pachitariu 2025).

#### Classification Panel

Users can define cell classes, assign or remove mask labels directly through an interactive image, or apply a pre-trained DenseNet classifier (Huang et al. 2017).

Once images are annotated with segmentation and classification data, the dataset can be used to train ML models to automate the process in the future and to create visualizations comparing the distribution of computed cell properties within/between classes. The image dataset can be downloaded with optional annotations, including overlaid masks colour-coded by class and cell type counts. Additionally, the user can download tabulated counts of cell types in each image.

### Mycol Workflow: Fine tuning ML models for pipeline automation

On the *Fine-Tune Models* page (**Supp. Fig. 3**), annotated datasets can be used to train new Cellpose (segmentation) or DenseNet (classification) models without coding expertise. Users may optionally adjust training parameters, although default settings have been selected to ensure robust performance across a wide range of tasks. On completion of training, Mycol generates a set of diagnostic plots to help the user evaluate fine-tuned model performance relative to the original model.

#### Cellpose fine-tuning diagnostic plots

training and validation loss across epochs, intersection-over-union (IoU) scores for predicted masks on a subsample of images for both the base and fine-tuned models, true-versus-predicted cell counts for both models on the same subsample.

#### DenseNet fine-tuning diagnostic plots: training and validation loss across epochs; accuracy, precision, and F1 scores for the fine-tuned model; and a confusion matrix to visualise class-specific performance

Trained models become immediately available for automated image analysis within the *Annotate Images* page. They can also be downloaded, along with the corresponding training histories, validation metrics, and diagnostic plots, for external use or for loading into a future app session. An example notebook is provided in the GitHub repository to help users replicate the image preprocessing when deploying trained models externally.

### Mycol Workflow: Visualizing and comparing morphological phenotypes

The *Cell Attributes* page enables users to explore a wide range of automatically computed descriptors for each segmented cell and to compare these features across different phenotypic classes (**Supp. Fig. 4**). Mycol uses established python packages to calculate commonly used morphological metrics, including measures of cell size, shape, elongation, compactness, and boundary irregularity (**Supp. Table 1**), providing users with quantitative summaries that can support tasks such as identifying morphological differences between conditions, validating classification results, or detecting outlier or low-quality segmentations. In-app explanations visualise what each metric described.

Users can visualise the distribution of these descriptors through class-wise comparisons (violin or bar plots with optionally overlaid datapoints) and all plots and tabulated measurements can be exported for downstream statistical analysis (**Supp. Fig. 5**). Together, these tools make it straightforward to quantitatively compare cell morphologies directly within the app, without requiring external software or coding expertise.

### Mycol for quantifying *Fusarium oxysporum* germination events

*Fusarium oxysporum* is a widespread soil-borne fungal pathogen responsible for devastating vascular wilt diseases in economically important crops (Jackson et al. 2024). A key step in its life cycle, and in the establishment of infection, is spore germination, where dormant spores transition into actively growing hyphae that can invade plant tissues (Gordon 2017). As a result, it represents a promising target for developing antifungal treatments or biocontrol strategies (Ortiz et al. 2021). However, there is a lack of fast, reliable, and easy-to-use methodologies for assessing how candidate compounds affect fungal spore germination. The commonly used approaches are based on Optical Density (OD) measurements (Hameed et al. 2024) or the acquisition of images at the microscope (Yang and Heinsohn 2007). Both these methods have limitations: OD measurements provide population-level information on fungal growth, without discriminating any phenotypic effects within the population, while the acquisition of images at the microscope requires tedious and error-prone human cell-counting and spore discrimination for each picture. This creates a bottleneck for large-scale or high-throughput experiments aimed at identifying factors that inhibit or promote fungal growth. We therefore hypothesised that Mycol would be useful for improving time efficiency when counting germinated and ungerminated *Fusarium oxysporum* cells.

In this work, we generated two *Fusarium oxysporum* datasets using Mycol. Firstly, 652 microscopy images containing 17,243 *Fusarium oxysporum* spores (**Fig. 2a**) were annotated with segmentation masks defining cell boundaries (**Fig. 2b**). A Cellpose2 model (Pachitariu and Stringer 2022) was fine-tuned on this dataset for 500 epochs using a learning rate of 0.1, weight decay of 0.0001, and batch size of 32 (**Supp. Fig. 6**); followed by hyperparameter optimisation to further improve segmentation performance (**Supp. Table 2**). The final model achieved an R^2^ of 0.981 when correlated with ground truth total cell counts in test set images (**Fig. 2c**). For the second dataset, 2,383 cell masks were labelled as germinated (944) or ungerminated (1439), and split 80:20 train:test for Densenet fine-tuning, achieving 95% accuracy, 95% precision, 95% recall and 94% F1 score (**Fig. 2d, Supp. Fig. 7**).

**Figure 2.**
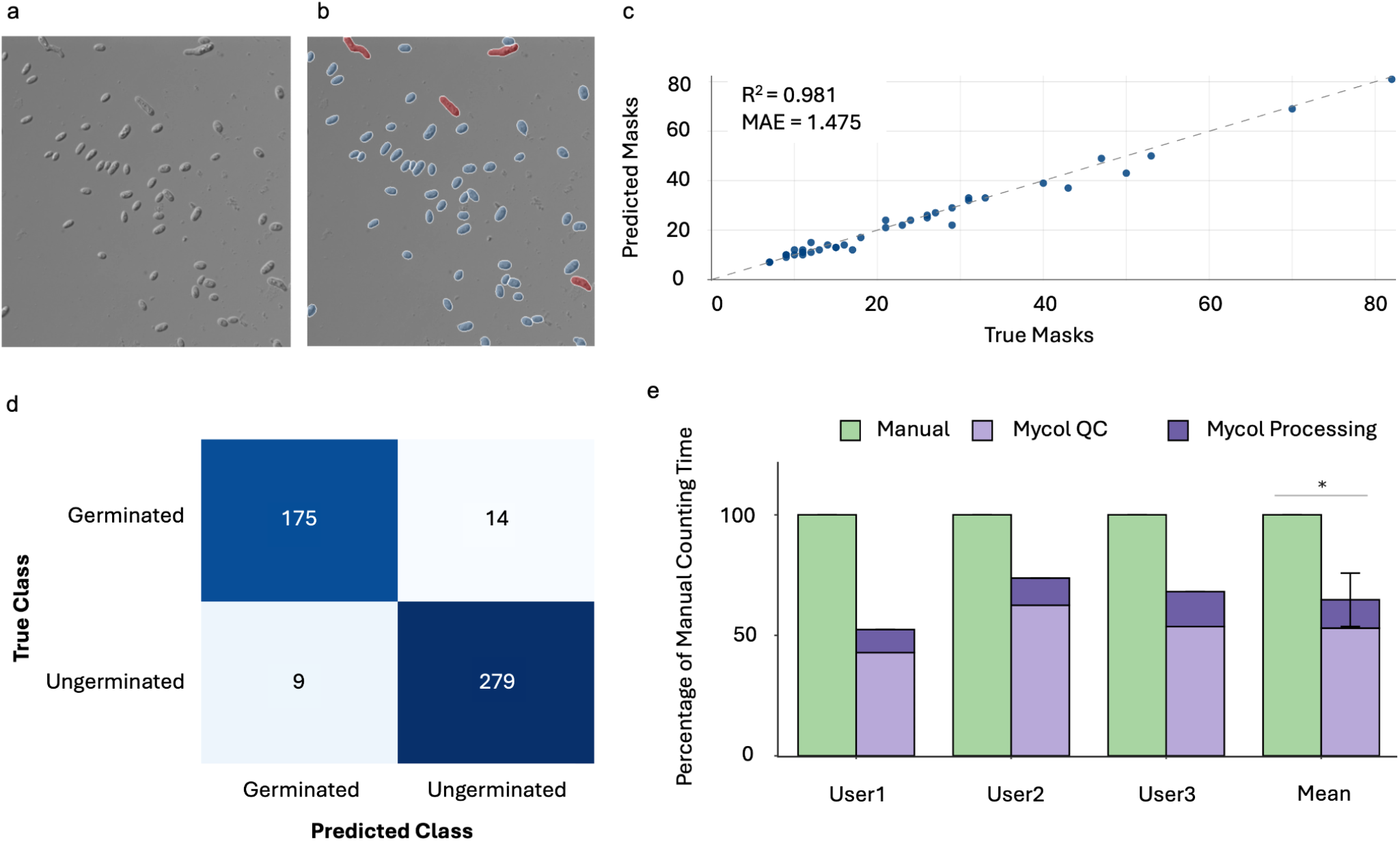
Applying Mycol to improve Fusarium oxysporum spore germination counting efficiency. **a**) representative microscopy image showing Fusarium oxysporum spores. **b**) same image in Mycol with cells highlighted with masks and classified as germinated (red) or ungerminated (blue). **c**) True cell count in Cyto2 fine-tuning test set images vs. predicted cell count by fine-tuned model. R^2^ and mean absolute error (MAE) are shown in the top left. **d**) Confusion matrix for fine-tuned Densenet model classifying cells as germinated or ungerminated in the test set. **e**) Time taken for three experimentalists to count 120 images of germinated and ungerminated Fusarium oxysporum spores with Mycol (Mycol QC) and without Mycol (Manual). ‘Mycol Processing’ indicates the time taken to run the segmentation and classification models (2 CPU cores and 2.7 GB RAM); Mycol QC indicates the time taken to Quality Control the automated predictions.

Time efficiency gains from Mycol were then benchmarked by three independent users counting *Fusarium oxysporum* germinated and ungerminated spores in two sets of 120 images. On average, the task took 35% less time to complete with Mycol than via manual cell counting (**Fig. 2e**), without compromising on consistency between the different experimentalists (**Supp. Fig 8**).

### Mycol for characterising mussel larvae development

The blue mussel (*Mytilus edulis*) is an edible marine bivalve mollusk of significant ecological and commercial importance. *M. edulis* is the most cultivated mollusk in European waters and global exports reached 300,000 tonnes in 2025 (*GLOBEFISH Quarterly Bivalves Analysis* 2026; Stępnicka et al. 2025). In addition, due to its sessile filter-feeding life strategy and wide distribution in temperate coastal habitats, *M. edulis* has become a model organism in marine pollution and climate change effects monitoring studies (Saidov and Kosevich 2021).

The standard practice for assessing healthy mussel larvae development, both for human consumption and assessment of their response to environmental and anthropogenic stressors, includes the visual discrimination of abnormalities in their D-shape stage of early development (His et al. 1997; Saidov and Kosevich 2021). Larvae are considered abnormal or malformed when they are not bilaterally symmetrical, have a protruding mantle or present a convex hinge in D-shaped larvae (His et al. 1997). This classification process is still performed manually in many cases, which is time consuming and prone to fatigue bias and subjective judgements as there are no universal quantitative conventions distinguishing malformation from natural variation (His et al. 1997; Saidov and Kosevich 2021; Thain 2013; His et al. 1999).

ML models in this field are limited and usually focused on model organisms from other taxons, such as the fruit fly *Drosophila melanogaster* and the fish *Danio rerio* (*GLOBEFISH Quarterly Bivalves Analysis* 2026; Stępnicka et al. 2025). We therefore hypothesised that Mycol could be used to identify and classify blue mussel larvae development abnormalities, making the process more objective and providing additional morphology data that could advance the understanding of *M. edulis* early development.

To evaluate the ability of Mycol to enhance analytical consistency and reporting fidelity, we analyzed 595 microscopy images of mussel larvae containing 885 larvae. Larvae were segmented using the cyto3 Cellpose model, which achieved robust performance without dataset-specific fine-tuning. Segmentation outputs were systematically reviewed and corrected where required, and 680 larvae were annotated as exhibiting normal or abnormal development (228 normal; 452 abnormal) (**Fig. 3a**). We next fine-tuned a DenseNet121 classifier on the curated dataset, yielding a model with classification accuracy and precision of 88% and an F1 score of 86% (**Fig. 3b; Supp. Fig. 9a, b**). By integrating segmentation, annotation, and classification within a unified and accessible framework, Mycol enables reproducible workflows and standardized reporting through the automated export of annotated images.

**Figure 3.**
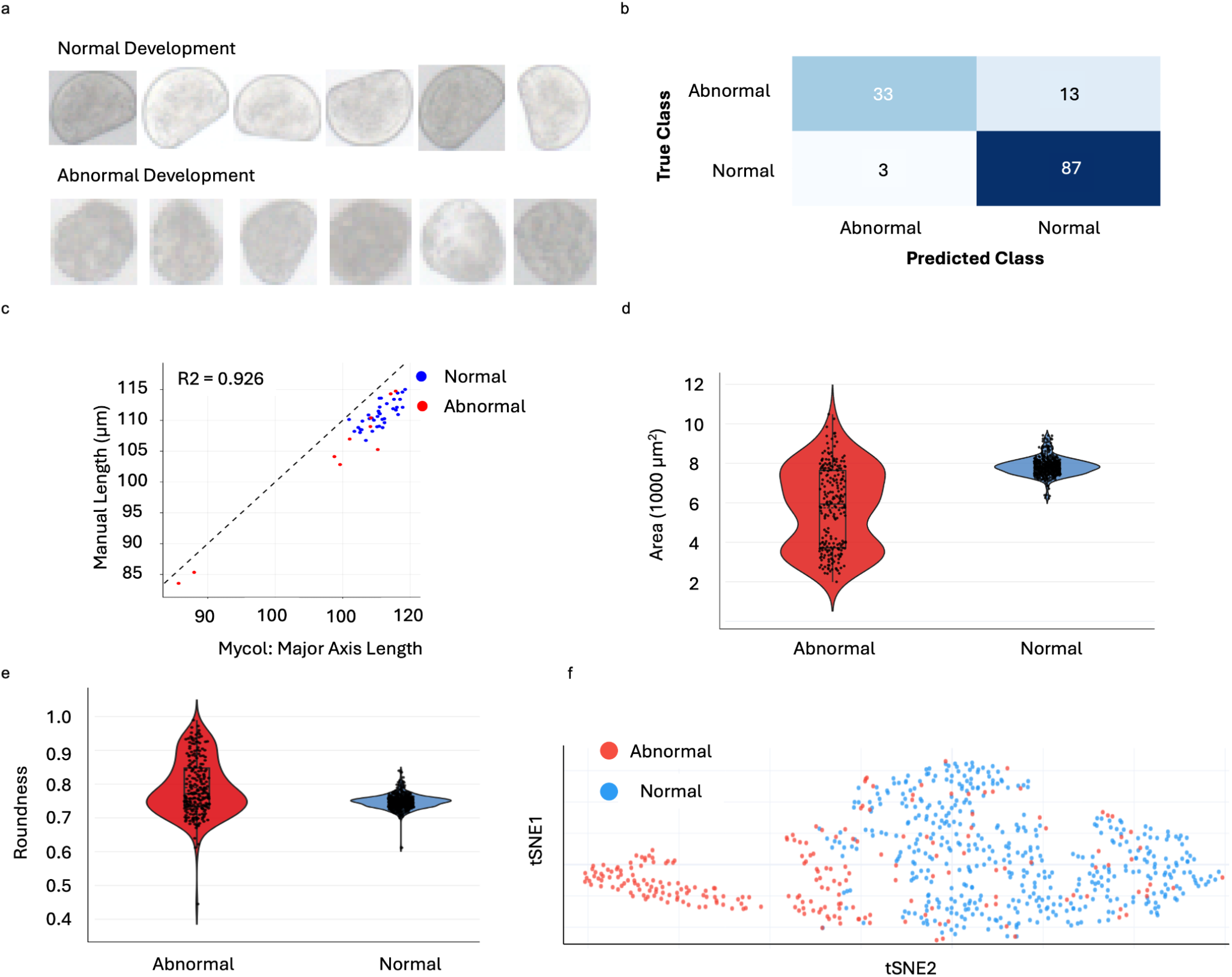
Applying Mycol to facilitate reproducibility and reporting in mussel larvae. **a**) Mussel larvae with normal (top) and abnormal (bottom) development, **b**) confusion matrix for DenseNet121 model fine-tuned to classify mussel larvae as normal or abnormal, **c**) correlation of 47 manually measured mussel larvae length (ImageJ) and the Major Axis Length parameter calculated in Mycol. **d**) distribution of area of normal and abnormal larvae, **e**) distribution of roundness of normal and abnormal larvae, **f**) tSNE visualisation of 14 morphological parameters calculated in Mycol for all labelled mussel larvae.

Quantitative analysis of the segmented larvae revealed close correlation (R^2^ = 0.926) between the major axis length parameter calculated in Mycol and manual length measurements (**Fig. 3c**), with a mean absolute difference of 4% (**Supp. Fig. 10**). Larvae length is an established parameter for assessing larvae development and we therefore propose Mycol as a convenient update to this manual methodology that is still in use in the field (His et al. 1999). Beyond this, distinct distributions of morphological parameters between normal and abnormal groups are also visible. In particular, abnormally developed larvae were generally smaller and more circular than normal larvae, although they also showed a much wider range of morphologies across the population (**Fig. 3d, e**). Notably, area measurements exhibited bimodal distributions within the abnormal group, suggesting the presence of at least two distinct modes or phases of developmental failure, which warrants further investigation (**Fig. 3d, e**). This observation was further supported by t-SNE visualisation of all 14 morphological parameters computed by Mycol, which separated abnormal larvae into two subpopulations: one distinct from normal larvae and another overlapping with them (**Fig. 3f**). These results highlight how automated feature extraction and visualization in Mycol can uncover biological patterns that may be overlooked in manual analysis.

## Conclusion

We developed Mycol to provide a flexible and accessible platform that unifies existing machine learning models within a single interface, reducing the expertise, time, and resources typically required for training, deployment, and quality control in cell image analysis. By combining intuitive annotation tools with efficient human-in-the-loop correction for both model training and prediction, Mycol addresses a persistent limitation of machine learning, its inability to achieve uniformly high accuracy across diverse tasks, thereby enabling consistently high-quality image annotations. By integrating annotation, correction, automated analysis, and interactive visualization into a single environment, Mycol lowers key barriers to adoption for non-specialists while remaining powerful for expert users. It streamlines small-to medium-scale workflows such as cell counting, image annotation, and phenotypic comparison, and provides an efficient framework for model development and validation. We demonstrate these capabilities by generating segmentation and classification datasets and fine-tuned models for two diverse organisms, *Fusarium oxysporum* and *Mytilus edulis*. In *F. oxysporum* germination assays, Mycol reduces processing time by approximately 35% relative to existing protocols, while in *M. edulis* development analysis, it enables accessible extraction of quantitative morphological data, which will facilitate future insights into the biological processes.

## Methods

### *F. oxysporum* sample preparation

Freshly obtained microconidia (1 × 10^9^) were washed with sterile double-distilled water, transferred to Germination media (Vitale et al. 2019), or Potato dextrose broth (PDB) to obtain the desired spore suspensions and incubated for 13 h to 15 h at 28 °C and 170 r.p.m.

### Mussel larvae sample preparation

Adult blue mussels (*M. edulis*) were kept in 50 L buckets with seawater and subsequently exposed to 10 - 20 minutes cycles of cold (10 °C) and temperate (20 °C) water to induce their spawning. Once spawned, their eggs and sperm were pooled and co-incubated for 72 h to facilitate fertilization. The resulting larvae were fixed with 800 µL of 4 % formalin and subsequently observed under an optical microscope.

### Microscopy image analysis

*F. oxysporum* images were collected with bright-field imaging with a Zeiss Axio Imager M2 microscope equipped with a Photometrics Evolve EMCCD camera, using a 40x oil objective. Mussel larvae images were collected with a Zeiss Axioscope 7 coupled with an Axiocam 305 color camera, using a 10x objective. Maximum axis length, defined as the longest distance parallel to the hinge (His et al. 1999), was measured with Image J (Rasband, W.S., ImageJ, U. S. National Institutes of Health, Bethesda, Maryland, USA, https://imagej.nih.gov/ij/, 1997–2018) by drawing a line with the Measure tool.

### Comparison between manual and tool-assisted counting

240 experimental images of *F. oxysporum* microconidia exposed to different media were equally split into two sets containing 120 pictures each. To avoid bias, both folders contained a comparable number of total cells and the same number of pictures of *F. oxysporum* exposed to each medium condition. The two folders were assigned to three PhD students with previous expertise in the quantification of *F. oxysporum* germination. The first folder was quantified manually, while the second was quantified using the tool. The three users reported the time spent with the two approaches, and in the case of the tool-assisted method, they also reported the time required to upload and analyse the 120 pictures to the Mycol app.

### tSNE Analysis

680 segmented *M. edulis* larvae annotated as normal (n = 452) or abnormal (n = 228) were described by twelve morphological descriptors calculated in Mycol (**Supp. Table 1**). Features were standardised to zero mean and unit variance i(sklearn.preprocessing.StandardScaler) prior to embedding. Two-dimensional t-SNE coordinates were computed with scikit-learn’s TSNE (perplexity = 30, 1000 iterations, random_state = 42).

### CRediT

S.A.B and G.S conceived the study. S.A.B, H.W and M.S wrote the software. G.S, A.P. and R. LM. acquired and analyzed *F. oxysporum* images. O. NS. acquired and analyzed *Mytilus edulis* images. S.A.B, G. S. and O. NS. wrote the manuscript. E. D. J., A. D. P. and C.G.A.R., secured funding, provided supervision and edited the manuscript.

## Data availability

Datasets and models for the two case studies are available for download from the Zenodo repository. Case study 1 (*F. oxysporum*): 10.5281/zenodo.20489251. Case study 2 (*M. edulis*): 10.5281/zenodo.20488331.

## Code availability

All code is open-source and available with installation instructions at https://github.com/biosustain/mycol, App overview and FAQ are available at https://biosustain.github.io/mycol/.

## Acknowledgements

We thank Ana Rodríguez López (University of Cordoba) for providing microscopy images of *Fusarium oxysporum*. We thank Kevin Ugwu (Roskilde University) and Sinja Rist (DTU Aqua) for their contributions to the successful spawning and maintenance of the larval cultures used in this study. Figures were created using *https://BioRender.com*.

