## Supplementary Information for "Mycol: A user-friendly app for automating analysis of microscopy images"

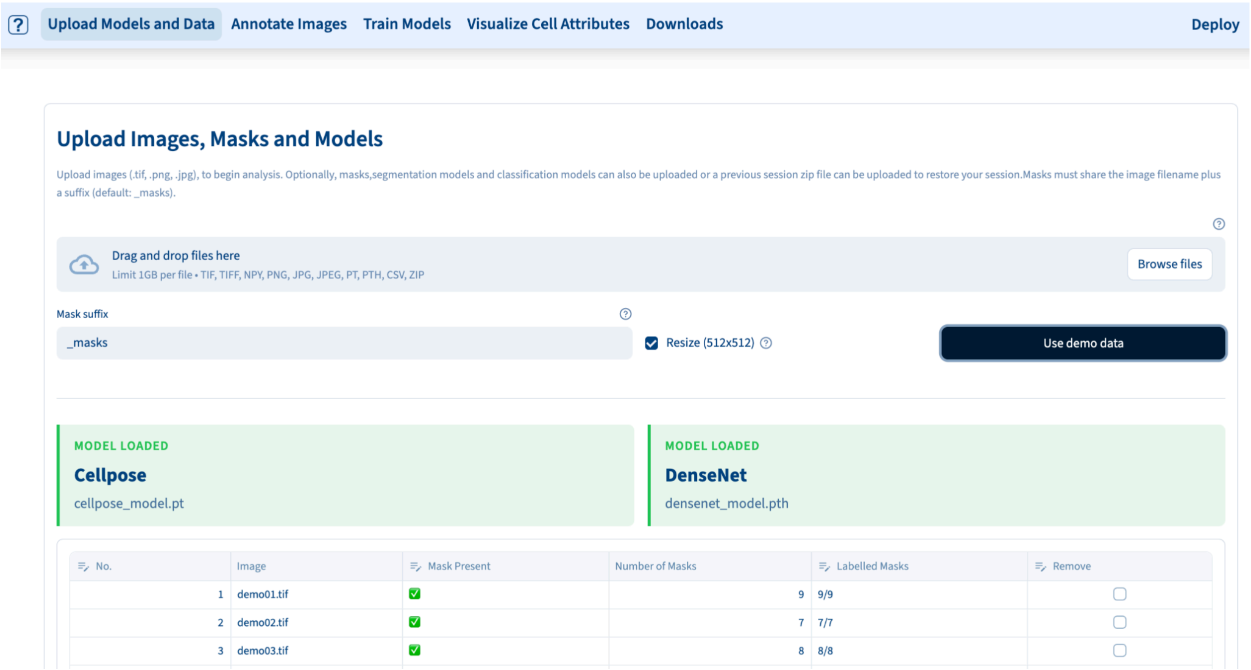


**Supplementary Figure 1. Mycol Uploads page layout**. Schematic showing the layout of options and readouts when uploading files into the Mycol application.

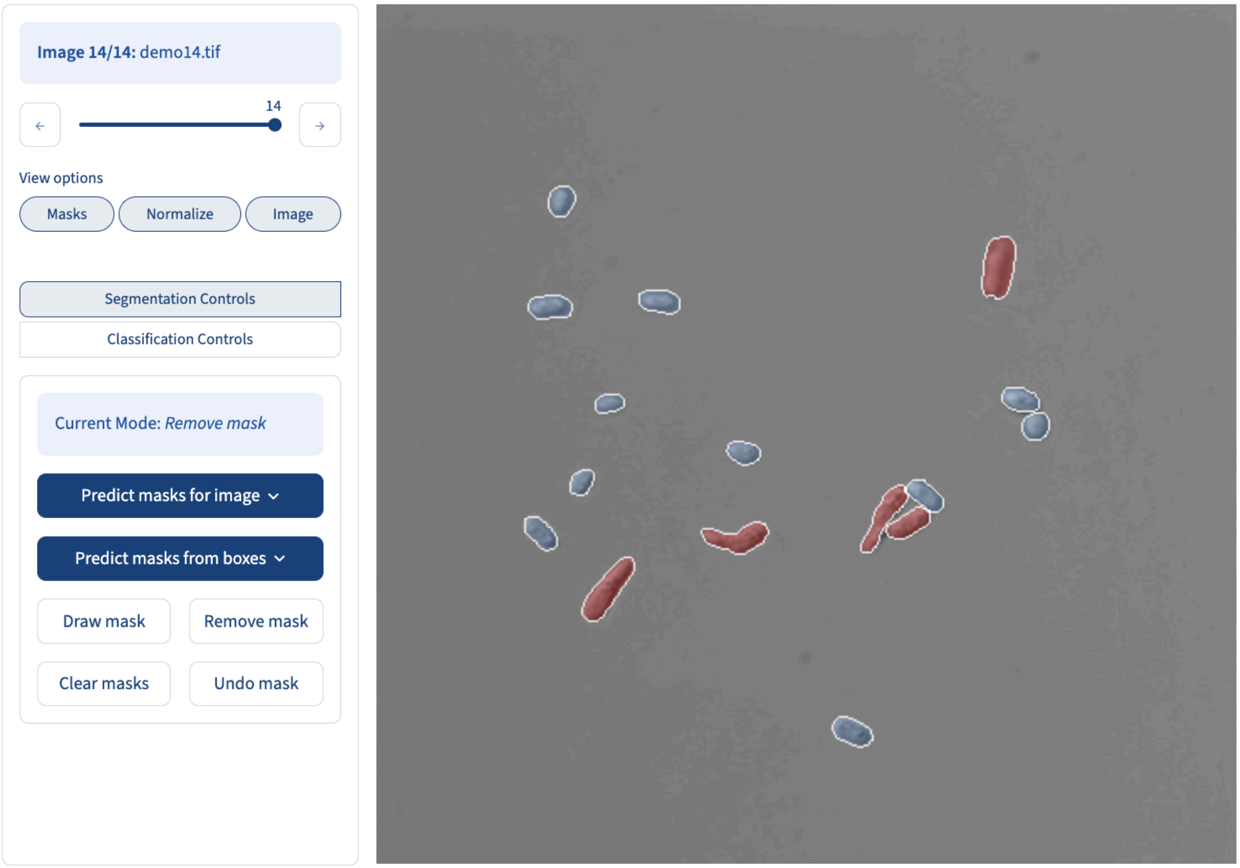


**Supplementary Figure 2. Mycol Annotations page layout**. Schematic showing the layout of options and readouts when segmenting and classifying cells in uploaded images.

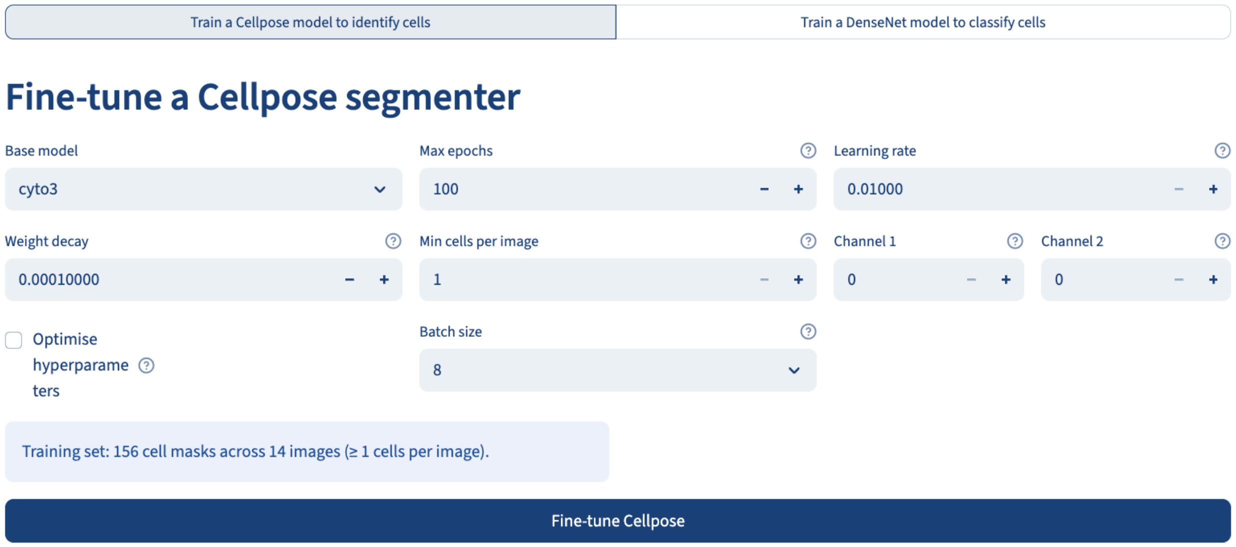


**Supplementary Figure 3. Mycol Fine Tuning page layout**. Schematic showing the layout of options and readouts when fine-tuning a Cellpose model in the Mycol application.

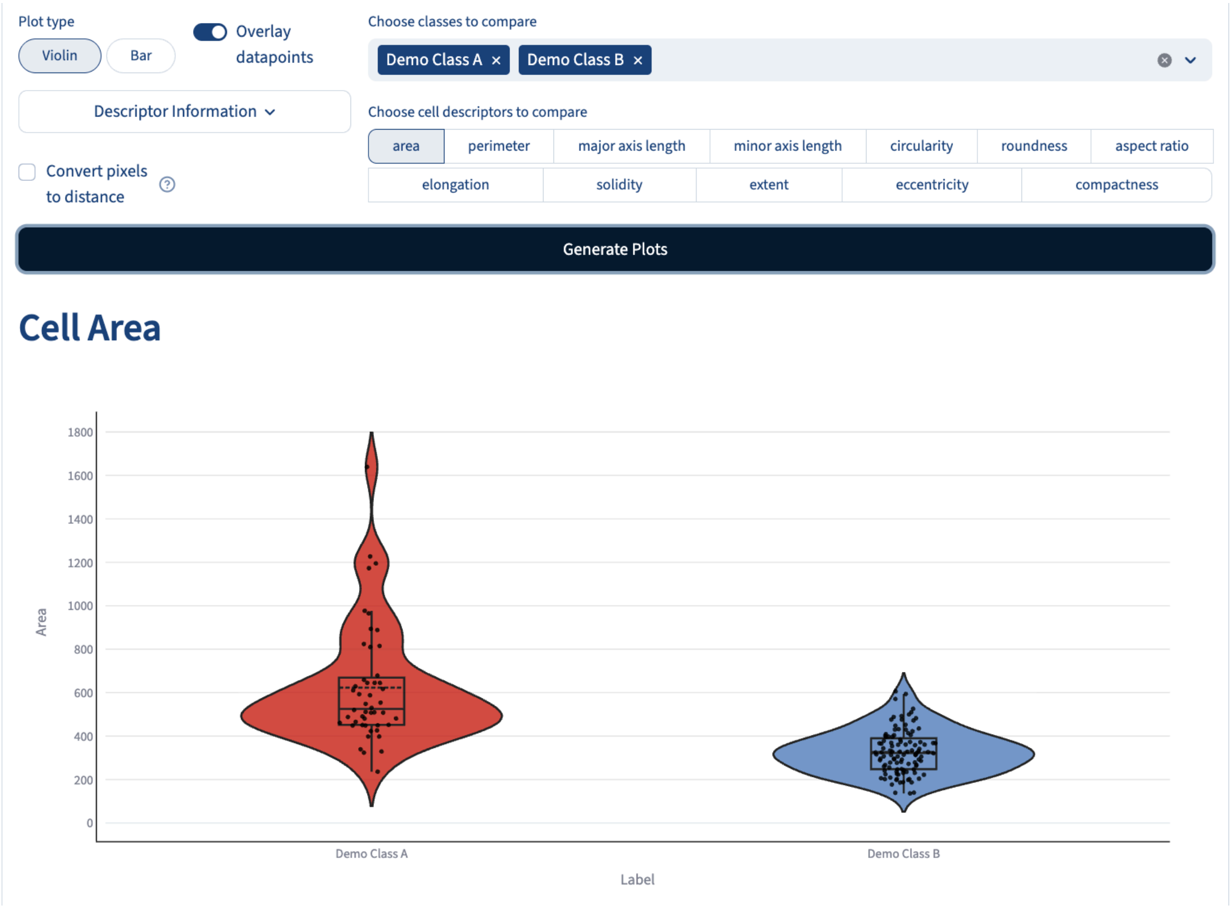


**Supplementary Figure 4. Mycol Cell Metrics page layout**. Schematic showing the layout of options and readouts when comparing morphological descriptors in the Mycol application.

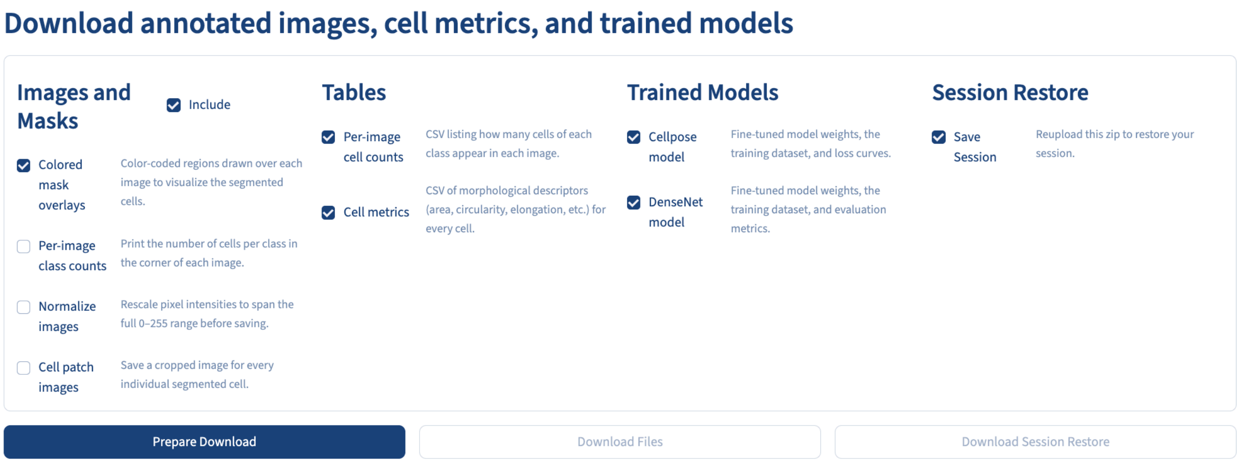


**Supplementary Figure 5. Mycol Downloads page layout**. Schematic showing the layout of options and readouts when datasets and analysis parameters from the Mycol application.

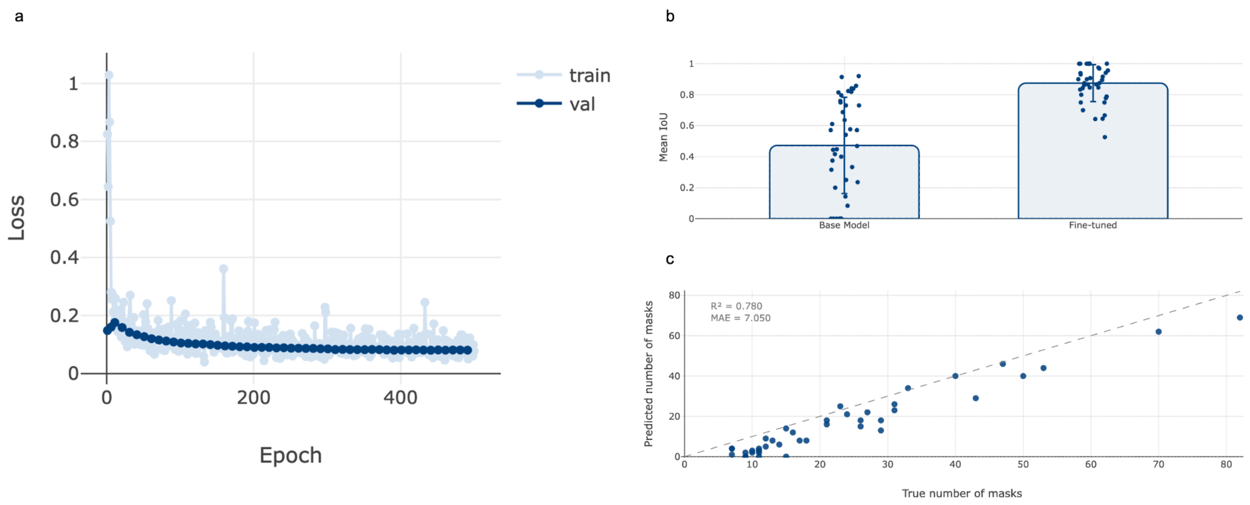


**Supplementary Figure 6. Fine tuning Cellpose2 to segment *Fusarium oxysporum* spores**. **a**) Per epoch model losses on train and test sets during fine-tuning. **b**) Mean fraction of masks in an image with IoU greater than 0.5 for the Cellpose2 model (base model) and the fine-tuned model (Fine-tuned) for the test set images. **c**) True per image cell counts for each image in the test set vs the Cellpose2 (pre-fine tuning) model.

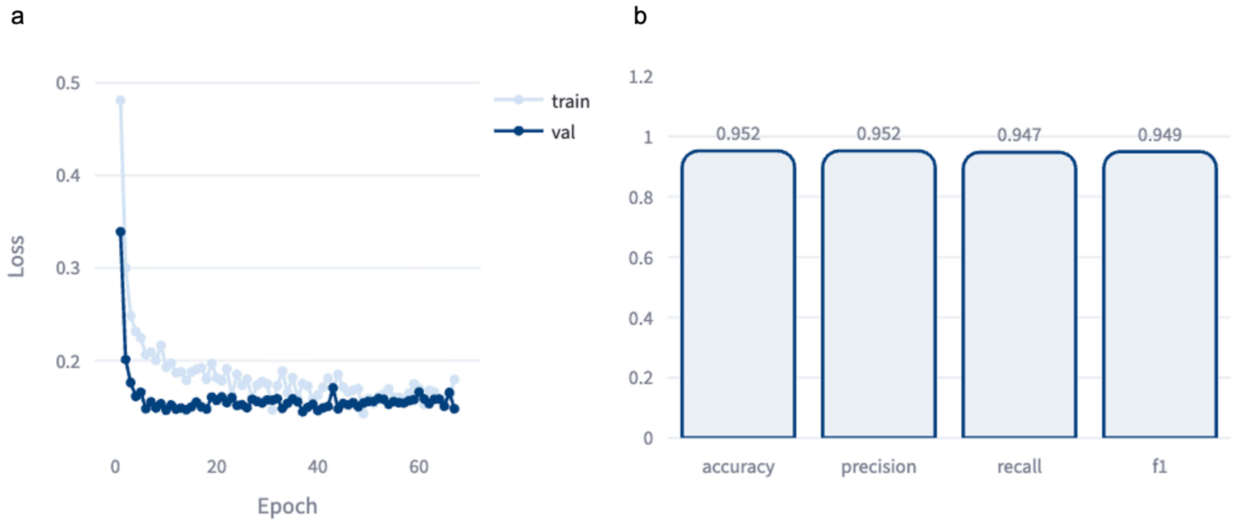


**Supplementary Figure 7. Fine tuning DenseNet121 to classify *Fusarium oxysporum* spores as germinated or ungerminated**. **a**) Per epoch model losses on train and test sets during fine-tuning. **b**) Accuracy, precision, recall and f1 scores of the fine-tuned model on the test set.

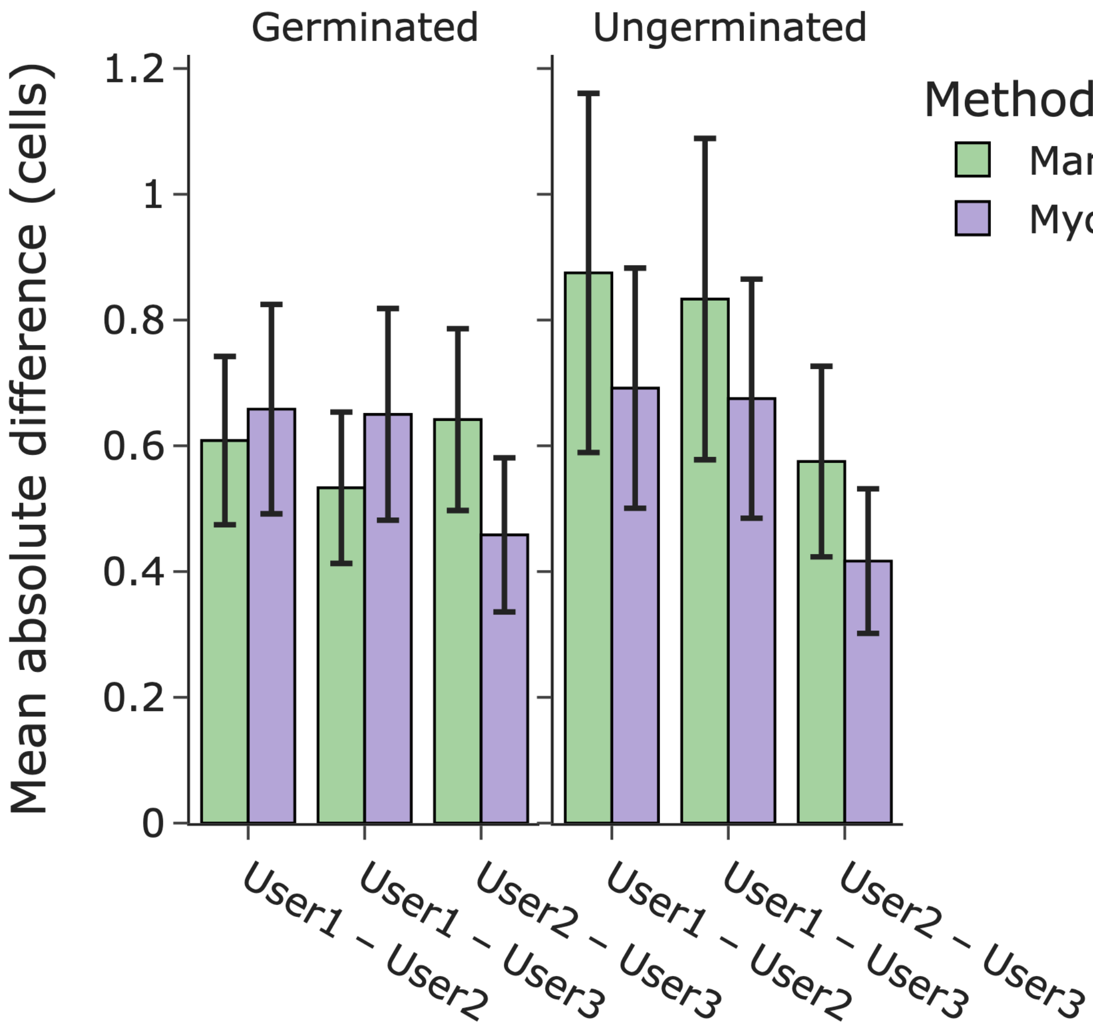


**Supplementary Figure 8. Comparison of germinated and ungerminated counts when counting manually or with Mycol assistance**. The mean absolute difference between cells count per image for germinated and ungerminated cell counts between three users. Plotted with 95% confidence limits.

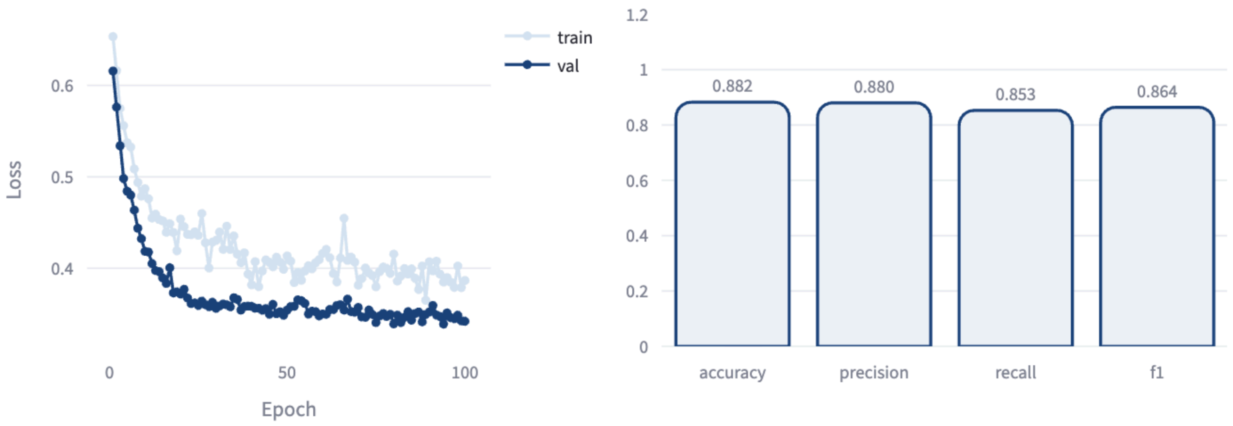


**Supplementary Figure 9. Fine tuning DenseNet121 to classify Mussel larvae as abnormal or normal**. **a**) Per epoch model losses on train and test sets during fine-tuning. **b**) Accuracy, precision, recall and f1 scores of the fine-tuned model on the test set.

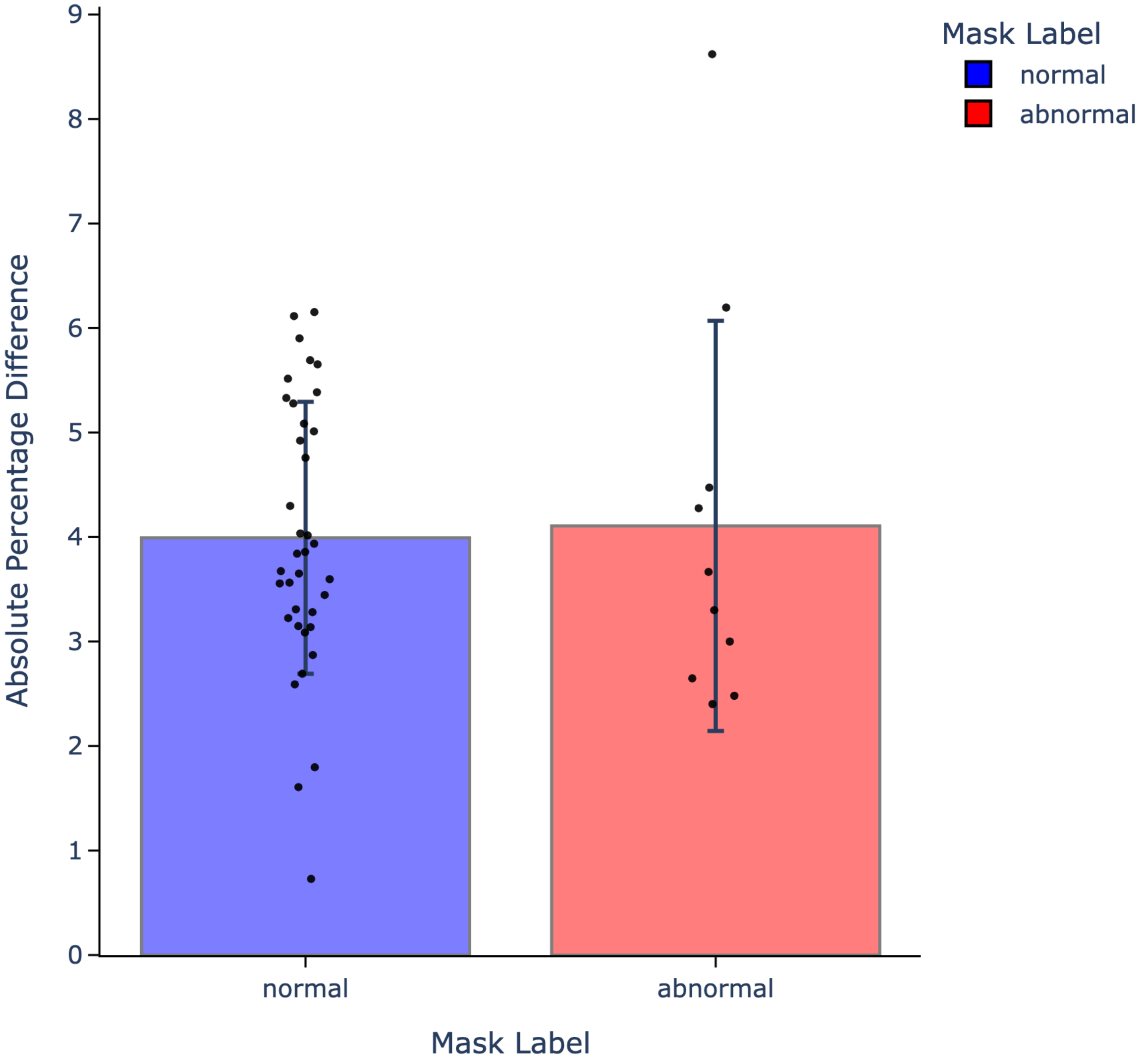


**Supplementary Figure 10. Mean absolute percentage difference between manually measured larvae length and Major Axis Length calculated from Mycol**. Mean values are shown for 47 measurements of normal (blue) and abnormal larvae (red) with standard deviation. Individual data points are overlaid in black.

**Supplementary Table 1. Calculated morphological descriptors returned by Mycol.**

| Property | Origin | Function |
|---|---|---|
| Area (A) | scikit-image | skimage.measure.regionprops: prop.area |
| Perimeter (P) | scikit-image | skimage.measure.regionprops: prop.perimeter |
| major axis length | scikit-image | skimage.measure.regionprops: prop.major_axis_length |
| minor axis length | scikit-image | skimage.measure.regionprops: prop.minor_axis_length |
| solidity | scikit-image | skimage.measure.regionprops: prop.solidity |
| extent | scikit-image | skimage.measure.regionprops: prop.extent |
| eccentricity | scikit-image | skimage.measure.regionprops: prop.eccentricity |
| circularity | mycol | $4 \cdot \pi \cdot A / P^2$ |
| roundness | mycol | $4 \cdot A / (\pi \cdot \text{major axis}^2)$ |
| aspect ratio | mycol | major axis / minor axis |
| elongation | mycol | (major axis − minor axis) / (major axis + minor axis) |
| compactness | mycol | $P^2 / (4 \cdot \pi \cdot A)$ |

**Supplementary Table 2. Hyperparameter search for fine-tuned Cellpose2**. Cellpose hyperparameter search values with mean fraction of masks in an image with an intersection over union (IoU) greater than 0.5.

| cellprob | flow_threshold | niter | min_size | ap_iou_0.5 |
|---|---|---|---|---|
| 0.0 | 0.4 | 900 | 100 | 0.8747802972793579 |

| | | | | |
|---|---|---|---|---|
| 0.0 | 0.4 | 800 | 100 | 0.8747802972793579 |
| 0.2 | 0.4 | 1000 | 100 | 0.8698738217353821 |
| 0.0 | 0.6 | 500 | 150 | 0.8541355133056641 |
| 0.4 | 0.0 | 600 | 50 | 0.8527186512947083 |
| 0.2 | -0.2 | 700 | 50 | 0.8491517901420593 |
| 0.6 | 0.0 | 200 | 100 | 0.8455308675765991 |
| 0.2 | 0.0 | 700 | 0 | 0.8425501585006714 |
| 0.6 | 0.2 | 1000 | 0 | 0.8378584980964661 |
| -0.2 | 0.2 | 800 | 100 | 0.8324806094169617 |
| 0.0 | 0.2 | 500 | 150 | 0.8297139406204224 |
| 0.0 | 0.4 | 900 | 200 | 0.8226549029350281 |
| 0.4 | -0.2 | 300 | 200 | 0.7801471948623657 |
| 0.2 | 0.4 | 200 | 250 | 0.7166754603385925 |

| | | | | |
|---|---|---|---|---|
| 0.0 | 0.4 | 800 | 300 | 0.6395906209945679 |
| 0.0 | 0.6 | 800 | 300 | 0.6343086361885071 |
| -0.2 | 0.6 | 900 | 350 | 0.5361915826797485 |
| -0.2 | 0.6 | 1000 | 400 | 0.42915934324264526 |
| -0.2 | -0.2 | 100 | 450 | 0.3613477647304535 |
| 0.4 | 0.4 | 100 | 500 | 0.3001483082771301 |
